# Discovery of phage defense systems through component modularity networks

**DOI:** 10.1101/2025.09.30.679545

**Authors:** Daan F. van den Berg, Ana Rita Costa, Jelger Q. Esser, Aswin Muralidharan, Halewijn van den Bossche, Stan J.J. Brouns

## Abstract

Phage defense systems in bacteria exhibit high degrees of modularity, with sensing, signal transmission, and effector enzymes frequently being exchanged among phage defense gene clusters. In this study, we capitalized on this modularity to discover phage defense systems by searching for defense-associated modules in new gene contexts. This approach revealed a large and interconnected network of modular components distributed across diverse gene clusters. From over 500 candidate defense systems, we selected nine for experimental testing and validated three: Dionysus, a TerB-encoding system that disrupts early phage infection vesicle formation by Jumbo phages; Ophion, a Radical SAM-containing system that prevents the formation of the Jumbo phage nucleus; and Ambrosia, a tightly regulated RM-like system. Collectively, we demonstrate that leveraging the modular architecture of phage defense systems is an effective approach to their discovery.

## NTRODUCTION

Bacteriophages (phages) are viruses that infect and replicate within a bacterial host^1^. To defend themselves against these viral threats, bacteria evolved a diverse repertoire of phage defense systems. These systems detect the infecting phage and trigger immune responses that range from cleaving the invading phage DNA to disrupting key metabolic processes, such as ATP and NAD^+^ metabolism, thereby preventing phage propagation^2^. In response, phages evolved counter-defenses, prompting further bacterial adaptations^3^. Over time, this ongoing evolutionary race has generated remarkable diversity among bacterial anti-phage strategies^2^.

It has become increasingly evident that many defense systems are built from modular components. This modularity is exemplified by phage defense families which comprise multiple subtypes that are unified by conserved core elements, such as the cyclase of CBASS, the ATPase NsnB of Menschen, LmuB of Lamassu, YprA family helicase of ARMADA, and the protease-nuclease pair of Canu^4–8^. Apart from shared genes, many systems have also interchanged functional domains, creating novel architectures from a common pool of modular parts. For example, sirtuin (SIR2) domains appear in several phage defense families, including Thoeris, prokaryotic argonautes (pAgos), defense-associated sirtuins (DSRs), and Avs^6,9–13^. Other frequently repurposed domains include superfamily helicase (SF2), HNH endonuclease, CD-NTase associated protein 4 (Cap4), modified DNA rejection and restriction (Mrr), restriction endonuclease (REase), Toll/interleukin-1 receptor/resistance protein (TIR), Trypsin-like, Caspase, Metallo-β-lactamase (MBL), and purine nucleoside phosphorylase (PNP) domains^7,8,14,15^.

The modular exchange of these genes and domains provides a genetic breeding ground for the genesis and diversification of phage defense systems. Understanding these modular components that are shared in evolution is key to uncovering the full spectrum of bacterial antiviral mechanisms. In this study, we use this principle to identify candidate defenses by analysing gene clusters that combine at least one known defense-associated gene with different domain contexts. We focused on *Pseudomonas aeruginosa*, a species with a highly diverse phage defense repertoire^16–18^. This approach uncovered more than 500 candidate systems, nine of which were selected for experimental testing. Of these, we validated three: Dionysus, a TerB-containing system specifically targeting jumbo phages; Ophion, a Radical SAM (rSAM)-containing system also targeting jumbo phages; and Ambrosia, a restriction-modification (RM)-like system targeting *Pbunavirus* and *Mesyanzhinovviridae* phages. Our findings demonstrate the power of modular domain-based discovery approaches to expand the known diversity of phage defense mechanisms and offer new insights into the architecture of bacterial immunity.

## Results

### Modularity of phage defense system components

More than 95% of known phage defense systems in *P. aeruginosa* reside within regions of genomic plasticity (RGPs)^19^. To identify previously uncharacterized phage defense systems, we performed an extensive search for gene clusters residing in these RGPs across 541 *P. aeruginosa* genomes. This analysis identified 1040 conserved gene clusters (**Figure 1A**) of which 515 (40%) contained functional domains associated with known defense systems. The most prevalent defense-associated domains were P-loop-containing ATPases (10.6%) and methyltransferases (MTases, 3.8%) (**Table S1**). The modularity of gene clusters that contained phage defense-associated functional domains was further analyzed using a gene network approach to investigate shared genes, resulting in a large, interconnected network of phage defense-related clusters (**Figure 1B-D, Supplementary File 1**).

**Figure 1.**
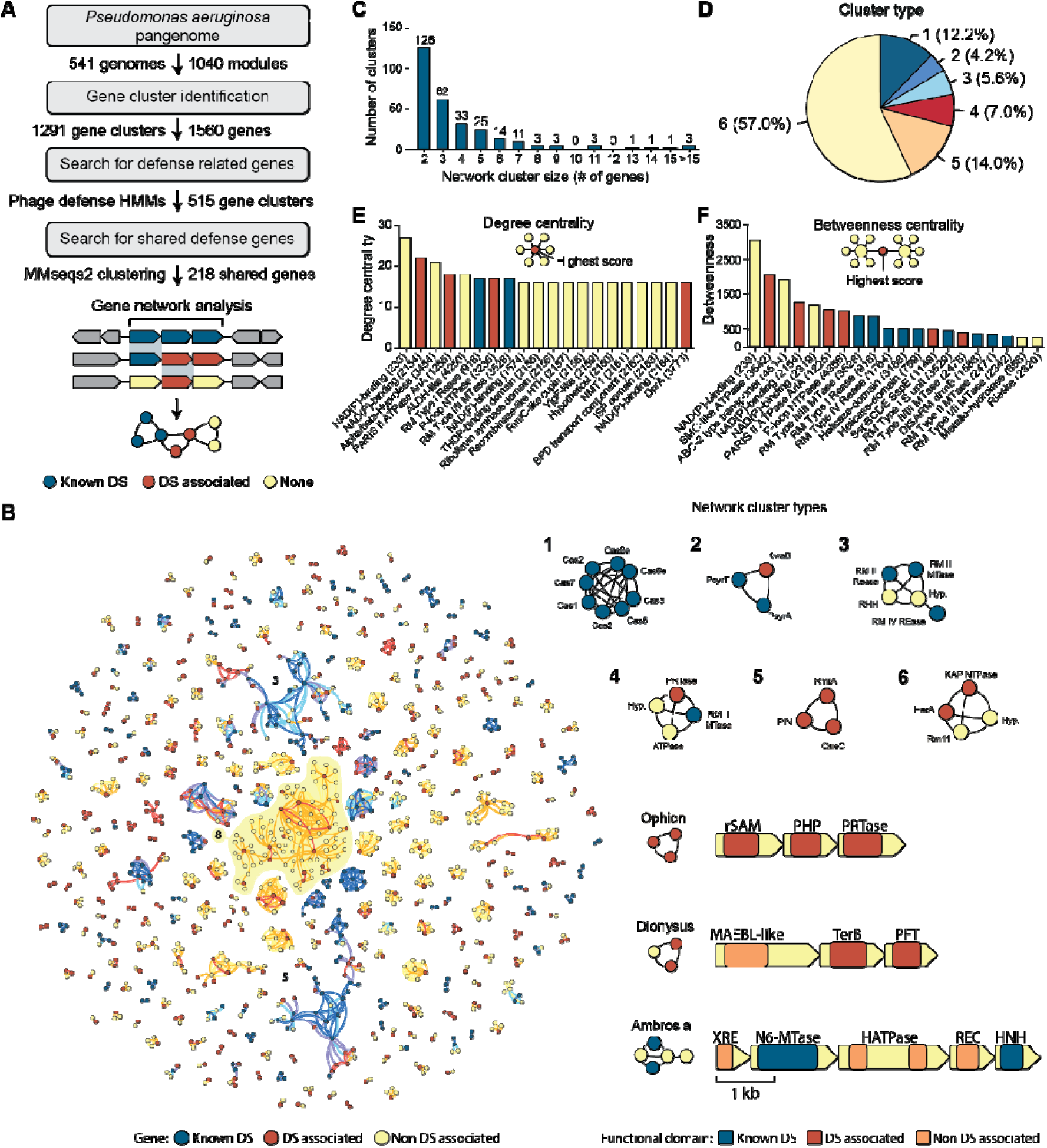
The gene network of phage defense system components in *Pseudomonas aeruginosa*. **(A)** Identification of gene clusters located in regions of genomic plasticity that contain functional domains previously associated with anti-phage activity. Gene clusters containing anti-phage functional domain were searched for shared genes, followed by generating a network of common components among these gene clusters. (**B**) Network of shared components between gene clusters containing anti-phage functional domains. Six types of network clusters were observed, including: 1. Complete defense systems; 2. Complete defense systems with additional defense-associated genes; 3. Complete defense systems with additional genes not previously associated with defense; 4. Complete defense systems with additional genes both defense-associated and non-defense-associated; 5. Clusters containing defense-associated genes in new combinations; and 6. Clusters with both defense-associated and non-defense-associated genes. The defense system clusters on the bottom right show the architecture and functional domains of the validated defense systems in this study. Genes are drawn to scale, with the scale bar representing 1 kilo base pairs (kb). (**C**) The size distribution of each cluster within the network. (**D**) Pie chart depicts the frequency of each cluster type in the network. (**E**) Number of connections that a gene establishes within the network of gene clusters containing anti-phage components (degree centrality). A higher score corresponds to a gene that is found together with more genes, either by being present in a large gene cluster or by being shared among multiple gene clusters. (**F**) Number of bridges that a gene establishes between subnetworks (betweenness centrality). A higher score corresponds to a gene more frequently shared among different families of phage defense systems.

We then classified the identified clusters into six distinct categories: 1. Complete defense systems (12.2%); 2. Complete systems with additional defense-associated genes (4.2%); 3. Complete systems with additional genes not previously associated with defense (5.6%); 4. Complete systems with additional genes both defense-associated and non-defense-associated (7.0%); 5. Clusters containing defense-associated genes in new combinations (57%); and 6. Clusters with both defense and non-defense genes (14%). Importantly, nearly half of the gene clusters (247/515; 48%) shared at least one gene with another cluster, allowing us to combine these into gene networks (**Figure 1B; Table S1**). Most of these networks are small, but three gene networks (networks 3, 5, and 8) each contain more than 15 genes (**Figure 1C; Table S1**). These larger networks consist of several gene clusters and may include novel or accessory elements relevant to phage defense. For instance, gene network 8, which is the largest network, spans 95 genes from 33 gene clusters and is largely composed of genes not previously associated with phage defense, aside from a few ATPases and STAND proteins. On the other hand, networks 3 and 5 are strongly enriched in known defense elements. Network 3 contains several type I and II restriction-modification (RM) system components, while network 5 includes genes that are part of BREX, DISARM, and Druantia systems. Although we did not further investigate these networks in this study, they offer starting points for future exploration of phage defense diversity and modularity.

To investigate the key contributors to defense network formation, we evaluated the gene connectivity within defense system families (degree centrality score: number of connections) and their role in linking distinct subnetworks (betweenness centrality score: bridge formation between subnetworks) (**Figure 1E,F; Table S1**). Interestingly, most genes with high number of connections within phage defense families (14 out of the top 20) have yet to be associated with phage defense, while known defense genes predominantly scored high in bridging phage defense subnetworks (15 out of the top 20) (**Figure 1E,F**).

For example, NAD(P)-binding protein (gene 233, **Table S1**) from gene network 3 is ranked the highest in both degree and betweenness centrality but has not previously been associated with phage defense. In contrast, other genes with high scores in both degree centrality and betweenness have been previously associated with phage defense, including MTases and REases of RM systems. Similar to what was observed for the largest gene networks 3 and 5, we found these to be commonly shared among RM subtypes as well as other RM-like systems^10,20,21^. Similarly, ATPases, such as SMC-like ATPase (gene 3542), PARIS ATPase AriA (gene 1225), and the ATP-Binding Cassette (ABC)-2 type transporter (gene 4514), scored highly in both categories. These ATPase domains are often phage-sensing components of defense systems, suggesting they might be shared among defense systems for phage specificity^22,23^.

These insights into the fluidity of the phage defense repertoire of *P. aeruginosa* suggested that we could utilize this gene network to discover additional phage defense systems. To test this, we selected nine gene clusters with shared defense components (clusters type 2-5) for experimental validation of phage defense and plasmid conjugation in *P. aeruginosa* strain PAO1^16^. Three gene clusters (33%), which we named Dionysus, Ophion, and Ambrosia (**Figure 1B**, **Table 1**), exhibited robust anti-phage activity and were further characterized, as discussed in detail below.

**Table 1.**
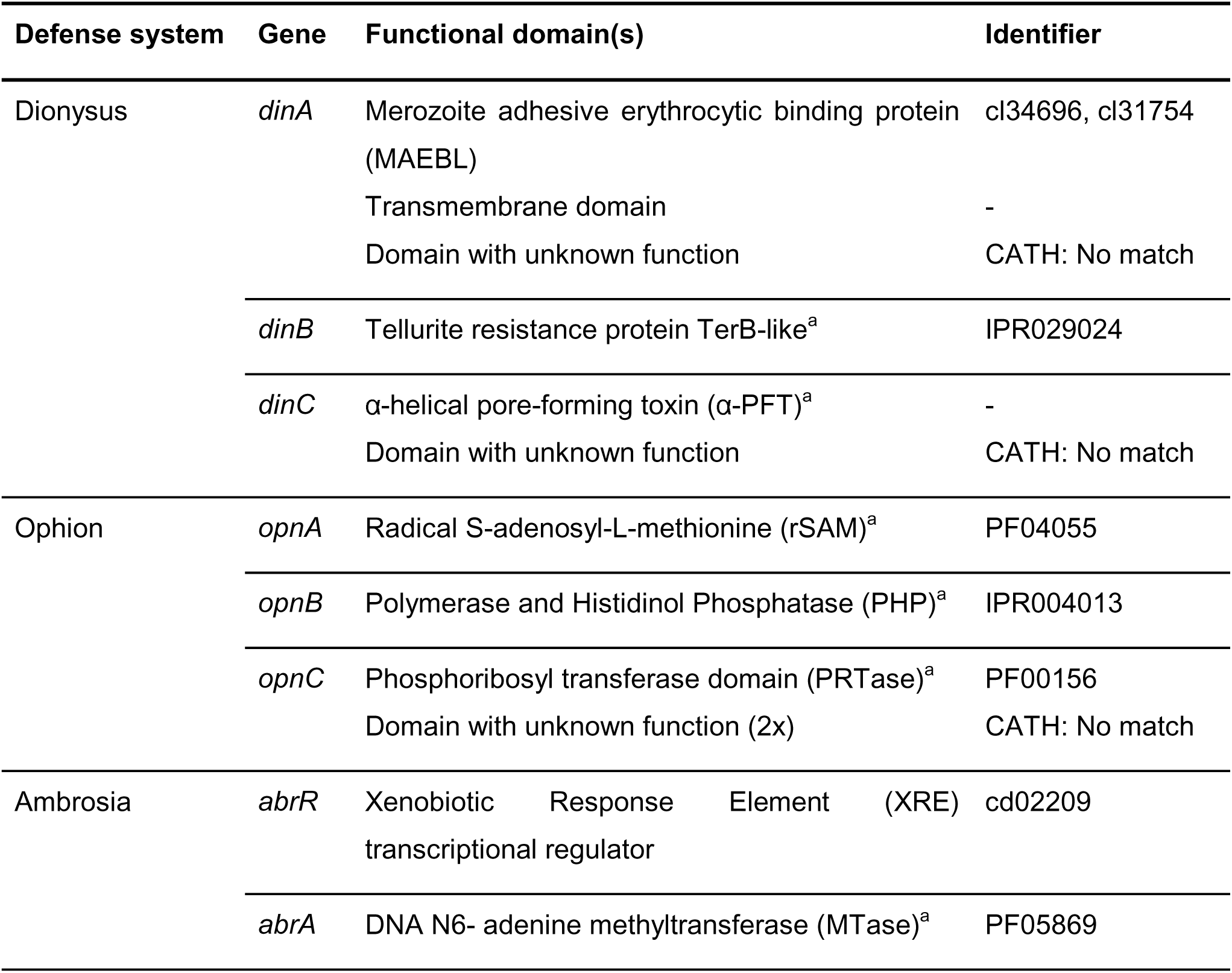
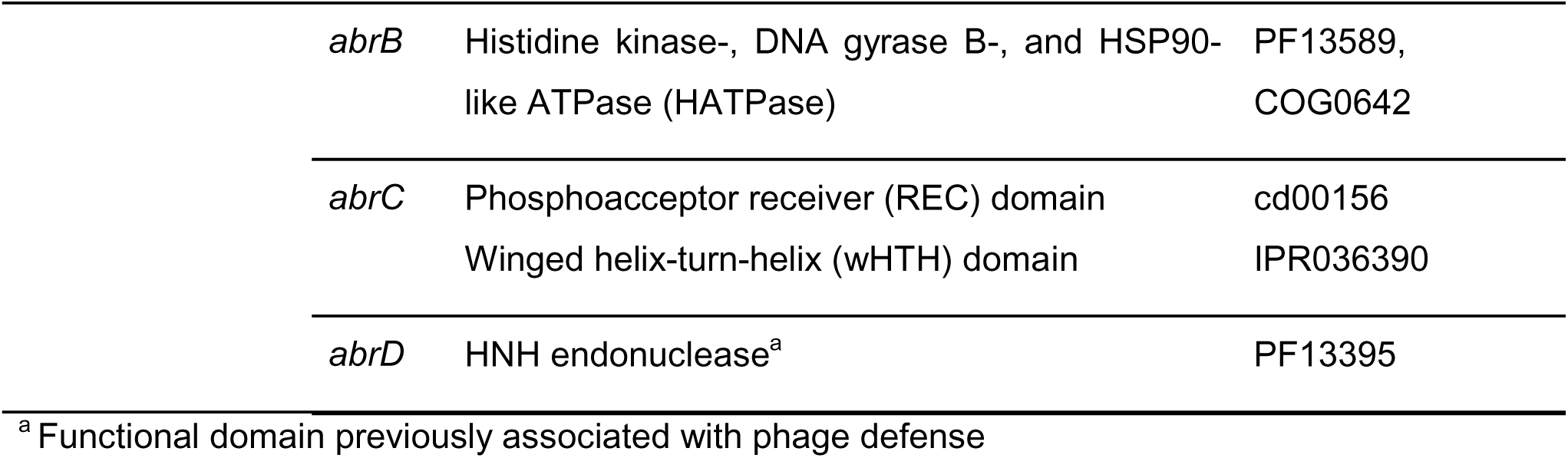
Genes and functional domains of the phage defense systems identified in this study

### Dionysus is a TerB-containing defense system with activity against *Phikzvirus* jumbo phages

Dionysus is named after the god associated with winemaking and the celebration of life. This defense system consists of three genes encoding: DinA, featuring a merozoite adhesive erythrocyte binding protein (MAEBL) and a transmembrane (TM) domain; DinB, a TerB-like protein; and DinC, a pore-forming toxin (PFT) with four TM domains (**Figure 2A**). Dionysus can be found in both Gamma-(312 instances) and Betaproteobacteria (49 instances) (**Figure S1A, Table S2**).

**Figure 2.**
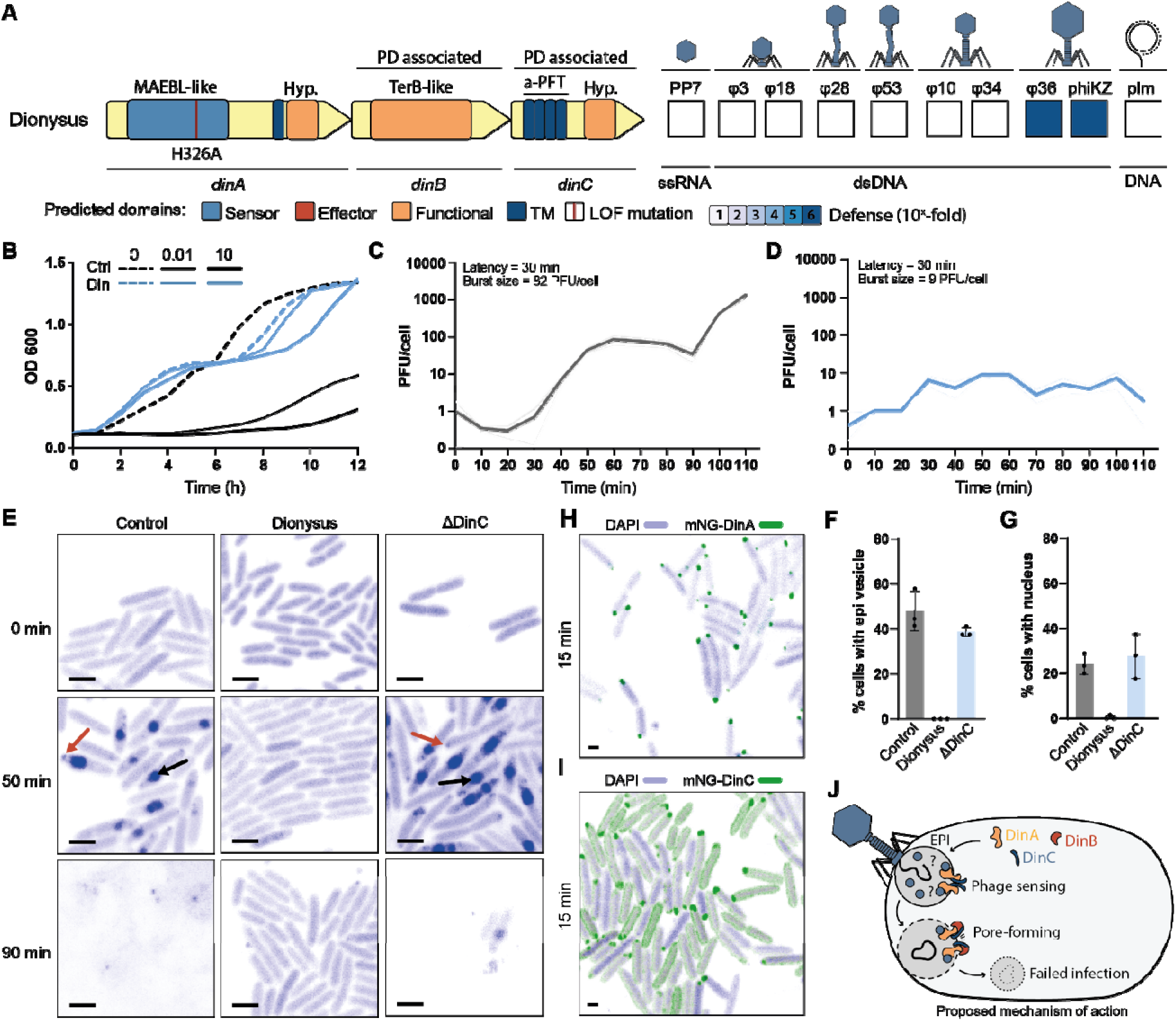
Dionysus is a jumbo phage specific defense system. **(A)** The functional domains of Dionysus, with loss of function mutation sites indicated as red lines. Genes are drawn to scale and indicated for their association with phage defense (PD associated). Dionysus was cloned with its native promoter into pUCP20 and introduced into *P. aeruginosa* strain PAO1, followed by assessing anti-plasmid activity (plm) by conjugation assays and anti-phage activity by efficiency of plating assays. The phage panel included *Fiersviridae* PP7, podophages *Autographiviridae* øPa3 and øPa18, siphophages *Casadabanvirus* øPa28 and *Mesyanzhinoviridae* øPa53, myophages *Pbunavirus* øPa10 and øPa34, and jumbo phages *Phikzvirus* øPa36 and phiKZ. **(B)** Effect of Dionysus on bacterial growth upon phage infection. *P. aeruginosa* strain PAO1 cells containing an empty plasmid (Ctrl) or Dionysus (Din) were infected with phage øPa36 at low (0.01) and high (10) multiplicity of infection, and their growth wa monitored for a period of 12h. **(C,D)** One-step phage growth curve of øPa36 infecting PAO1 with (C) empty plasmid and (D) Dionysus. **(E)** Confocal microscopy images during øPa36 infection (0, 50, and 90 min) of PAO1 with empty plasmid, Dionysus, and a knock-out of DinC (ΔDinC). 4′,6-diamidino-2-phenylindole (DAPI) was used to stain phage and host DNA. The scale bar is 1 μm in length. **(F,G)** The percentage of observed (F) EPI and (G) nuclei during øPa36 infection of control, Dionysus-expressing, and ΔDinC cells. (**H,I**) Confocal microscopy images during øPa36 infection (15 min) of Dionysus cells with (H) DinA-mNeonGreen and (I) DinC-mNeonGreen. The scale bar is 1 μm in length. (**J**) Working model for phage defense by Dionysus, which is hypothesized to form pores in the EPI vesicle upon phage infection, protecting against jumbo phages.

Dionysus provides robust protection against jumbo *Chimallivirus* phiKZ and øPa36, both in efficiency of plating (>10^6^-fold reduction) and liquid cultures (**Figure 2A,B**). The one-step growth curve of øPa36 in control and Dionysus cells revealed that the life cycle of the phage is severely compromised by the defense system, with a similar latency period but a significantly reduced burst size (82 ± 24 PFU/cell in control cells versus 9 ± 2 PFU/cell in Dionysus-expressing cells) (**Figure 2C,D**). All three proteins of Dionysus are essential for its protective capacity, as mutating a conserved amino acid within the MAEBL-like domain of DinA (H326A), and deletion of DinB or DinC, resulted in the partial (DinA H326A) or complete loss of anti-phage activity (**Figure S1B**). Lastly, *in silico* co-folding with AlphaFold 3 and further analysis using AlphaBridge predicts DinC to interact with the C-terminus of DinA to form a complex (1:1 stoichiometry; piCSi: 0.77), and DinB to form a homodimer (piCSi: 0.83) (**Figure S1C**).

Dionysus genes *dinB* and *dinC* encode domains that have previously been associated with phage defense. DinB has a TerB domain, which was originally identified in the tellurite resistance gene cassette and believed to be involved in the bacterial stress response^24,25^. More recently, TerB domains were also found in phage defense systems such as Bunzi (BnzA) and Shango (SngA)^6,17^ (**Figure S2D**). It is still unclear how TerB domain-containing phage defense proteins convey phage protection. It has been proposed that this could be related to TerB proteins having amphitropic characteristics^26^, meaning they can reversibly switch between the cytoplasm and membrane, depending on environmental signals or ligand binding. These previous observations suggest that DinB may be involved in membrane surveillance.

DinC contains a PFT domain, which has also been reported to act as the anti-phage effector in various defense systems, including CBASS, Pycsar, CRISPR-Cas, Retrons, and bGSDM^2,4,17,27,28^. PFTs often form multi-subunit pores with a hydrophilic interior in the cellular membrane to depolarize the bacterium and block phage propagation and can be broadly classified into two categories based on their secondary structure: α-PFTs, which form α-helical pores, and β-PFTs, which form β-barrel pores^29^. Inspection of the AlphaFold3 modelling of DinC revealed that it is an α-pore-forming toxin (α-PFT) consisting of multiple amphipathic α-helices, structurally similar to enterotoxins such as haemolysin BL (Hbl), non-haemolytic enterotoxin (Nhe), and cytolysin A (ClyA)^30^ (**Figure S1E**).

Dionysus gene *dinA* encodes an MAEBL domain, not previously linked to phage defense. This domain is poorly understood, apart from its role in *Plasmodium* species, where MAEBL is a transmembrane protein that binds erythrocyte receptor^31^. Therefore, DinA may act as a membrane-associated protein involved in detecting phage components during phage infection.

Based on the predicted function of DinA, DinB, and DinC, we hypothesized that Dionysus may convey phage protection by detecting membrane-associated processes of the jumbo phage through DinA and DinB, with the end result of DinC forming pores in the cellular membrane to cause cell death and abort phage infection. To investigate this hypothesis, we assessed membrane permeabilization using propidium iodide (PI) staining. We observed that at 90 min post infection by jumbo phage øPa36, the vast majority of Dionysus cells were not lysed, suggesting that Dionysus does not act through damaging of the cellular membrane (**Figure S1F**). This is further supported by quantification of cell survival at both one hour and two hours post infection, where no control cells survived after 2 hours, while Dionysus cells remained viable (**Figure S1G**). These results show that DinC does not form pores in the host membrane. We therefore hypothesized that DinC might act on the early phage infection (EPI) vesicles that jumbo phages produce to protect their DNA from host defense systems before transitioning into their nucleus-like compartment^32^. To explore this possibility, we performed confocal microscopy using 4′,6-diamidino-2-phenylindole (DAPI) to stain phage and host DNA, expecting that DNA foci appear at EPI vesicle and nucleus compartments. These compartments shield the phage DNA from DNA-targeting host defenses^33^. This experiment revealed no EPI vesicles or nuclei in cells expressing Dionysus, while both compartments could be observed in control (empty plasmid) and ΔDinC mutant cells (**Figure 2E-G; S1H**). At 50 minutes post-infection, EPI vesicles were present in almost half of the control and ΔDinC-expressing cells (48% and 39%, respectively), while none could be detected for Dionysus-expressing cells (**Figure 2F**). Moreover, no nuclei were detected in Dionysus cells, while 24% control and 28% ΔDinC cells contained nuclei (**Figure 2G**). These results were not affected by differences in adsorption, which was found to be identical between control and Dionysus-expressing cells (**Figure S1I**). To further investigate the potential interaction of Dionysus with the EPI vesicles of the Jumbo phage, we labelled either DinA or DinC with mNeonGreen and observed that, 15 min post infection, both proteins form foci where the EPI vesicles are formed, suggesting possible interactions between these proteins and further supporting the hypothesis of early activity of Dionysus on the EPI vesicle (**Figure 2H,I; Figure S1J**). Together, these results show that Dionysus specifically acts on the phage-derived EPI vesicle upon phage detection, preventing progression of jumbo phage infection at the first stage in its infection cycle (**Figure 2J**).

### Ophion is a radical SAM-containing anti-jumbo phage defense system proposed to act via nucleic-acid interfering mechanisms

Ophion, named after a primordial serpent from Greek mythology, is a three gene system encoding a radical S-adenosyl-L-methionine (rSAM) enzyme (OpnA), a polymerase-and histidinol-phosphatase (PHP, OpnB), and a phosphoribosyl transferase (PRTase, OpnC) (**Figure 3A**). Distributed across diverse Gammaproteobacteria (**Figure S2A; Table S2**), Ophion shows strong protection against jumbo phages øPa36 and PhiKZ, reducing phage infectivity by at least 10^6^-fold in EOP assays and blocking phage propagation in liquid cultures (**Figure 3A,B**). The one step growth curve of øPa36 in Ophion cells showed no detectable burst, indicating that the phage life cycle is completely disrupted (**Figure 3C**). Mutating conserved residues of the rSAM and PRTase domains revealed these to be essential for protection, with partial dependence on the PHP domain (**Figure S2B**).

**Figure 3.**
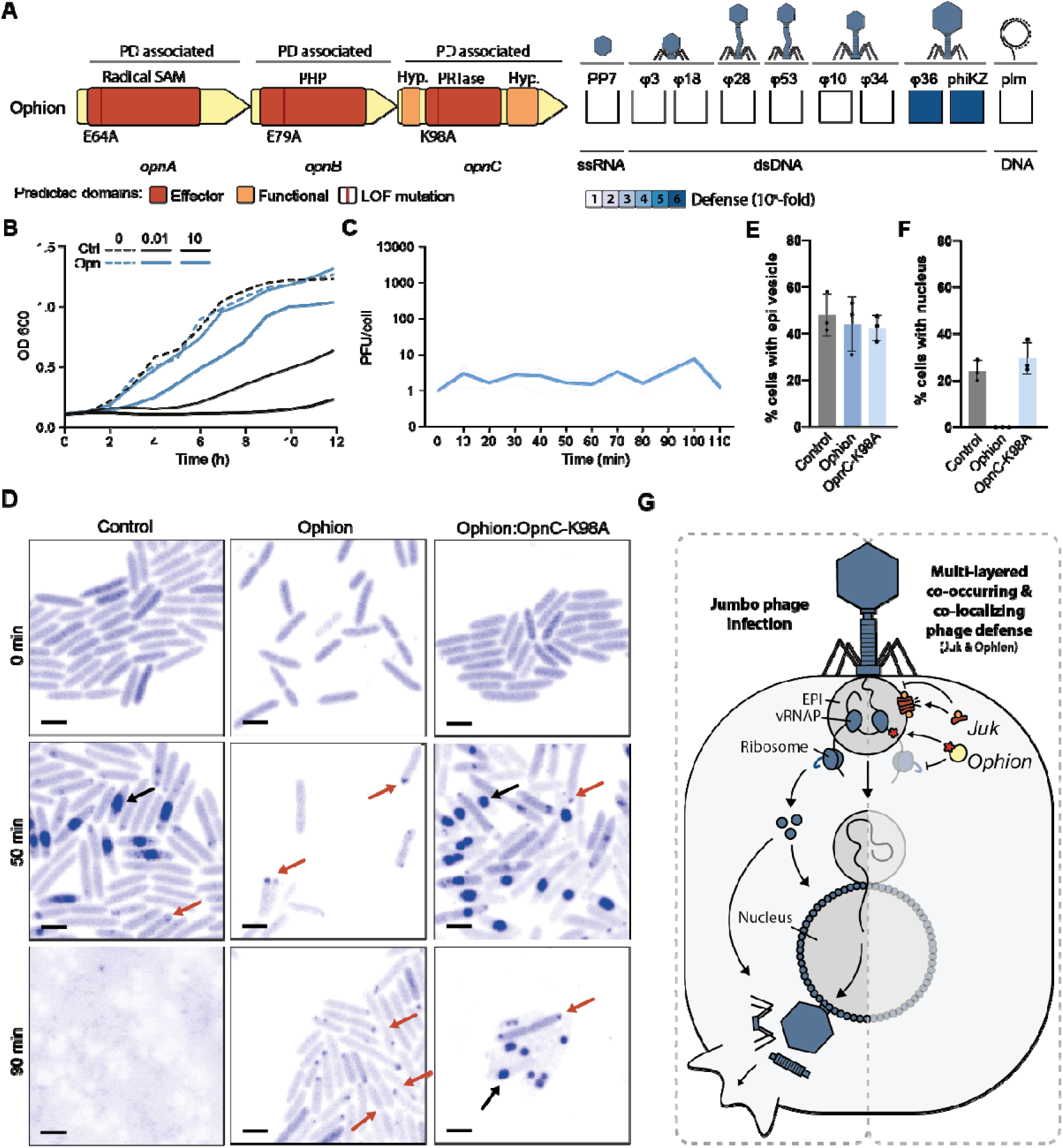
Ophion is a radical SAM-containing phage defense system proposed to act via nucleic-acid interfering mechanisms. **(A)** The functional domains of Ophion and the mutation sites (shown as red lines) that were tested. Genes are drawn to scale and indicated for their association with phage defense (PD associated). Ophion was assessed for anti-plasmid activity (plm) by conjugation assays and for anti-phage activity by efficiency of plating assays. The phage panel includes *Fiersviridae* PP7, podophage *Autographiviridae* øPa3 and øPa18, siphophage *Casadabanvirus* øPa28, siphophage *Mesyanzhinoviridae* øPa53, myophage *Pbunavirus* øPa10 and øPa34, and jumbo phages *Phikzvirus* øPa36 and phiKZ. **(B)** Effect of the defense system on bacterial growth upon phage infection. *P. aeruginosa* strain PAO1 cells containing an empty plasmid (Ctrl) or Ophion (Opn) were infected with phage øPa36 at low (0.01) and high (10) multiplicity of infection, and their growth was monitored for a period of 12h. **(C)** One-step growth curve of øPa36 infecting PAO1 with Ophion. **(D)** Confocal microscopy images during øPa36 infection (0, 50, and 90 min) of control (empty plasmid), Ophion, and OpnC mutant (K98A) cells. Both the bacterial and phage DNA were labeled using 4′,6-diamidino-2-phenylindole (DAPI). **(E,F)** The percentage of observed (E) EPI and (F) nuclei formed during øPa36 infection of control (empty plasmid), Ophion, and OpnC-K98A cells. (**G**) Proposed multilayered defense against jumbo phages. On the left side, jumbo phage progresses through its infection cycle, from formation of the EPI vesicle (early stage) to the transition into the nucleus stage, and the final assembly of the phage particles before lysing the host. On the right, jumbo phage infection is stalled at early stages when co-occurring and co-localizing jumbo phage defense systems Juk (targeting epi vesicle) and Ophion (likely targeting early phage transcripts) are present.

Each Ophion protein contains a domain associated with anti-phage mechanisms. For instance, the rSAM domain of OpnA is related to, but phylogenetically distinct from, those in prokaryotic Viperins (pViperins) (**Figure S2C**), which inhibit phage propagation by generating 3’-deoxy-3’,4’-didehydro-nucleotides (ddh-NTPs). These nucleotide analogs terminate RNA synthesis when incorporated into phage transcripts^34,35^. The PHP domain of OpnB is found in the ppl phage defense system and in family X DNA polymerases where it is involved in DNA proofreading as a metal-dependent exonuclease of the phosphate backbone, removing misincorporated nucleotides^12,36,37^. The PRTase domain of OpnC is known to play crucial roles in the biosynthesis of purine, pyrimidine, and pyridine nucleotides, as well as in purine and pyrimidine salvage^38^. PRTases were previously identified as the phage defense effector of the phage defense type III retron systems and as the toxin in the toxin-antitoxin phage defense systems ShosTA and PsyrTA^6,39^. These phage defense associated PRTases are phylogenetically related and located within the same clade of the PRTase family, suggesting a shared function (**Figure S2D**). Of these phage defense-associated PRTases, only the function of the ShosTA PRTase has been characterized, which disrupts the purine metabolism by producing nucleoside monophosphate, eventually disrupting DNA duplication of both the host and the phage^40^. The Ophion proteins are not predicted to interact with each other (**Figure S2E**).

Together, these observations indicate that each protein encoded by Ophion contains functional domains that are known to interfere with nucleotide modification and salvage pathways. To gain a broader understanding of Ophion’s potential mechanism of action, we set out to identify the stage of phage infection at which Ophion acts, using confocal microscopy to monitor øPa36 jumbo phage infection. In control cells and in cells expressing an OpnC mutant (K98A), the phage progressed normally through both early (EPI vesicle) and late (nucleus) stages. In contrast, in Ophion-expressing cells, øPa36 failed to develop nuclei (**Figure 3D-F**), indicating that infection is arrested early, before the translation of nucleus-forming components^41^. Given the nucleotide-modifying potential of the rSAM domain of OpnA and PRTase of OpnC^34^, it is possible that early phage transcription within the EPI vesicle is disrupted by incorporating transcription-halting nucleotides. Notably, Ophion does not seem to affect the host and does not trigger abortive infection (**Figure S2F**), implying a phage-specific mechanism.

Notably, we observed that Ophion strongly co-localizes and co-occurs with Jumbo phage defense system Juk^42^ in 98.2% of the cases (108 out of the 110 times Ophion is found in *P. aeruginosa*; Observed = 108; Expected = 23; Z-score = 18.25; Adjusted p-value = 0; Mean distance = 0 bp). Interestingly, Ophion and Juk convey protection against jumbo phages at different stages of their infection cycle. Specifically, Juk disrupts EPI vesicles which form early on in the infection, while Ophion affects the phage in a later stage by blocking the progression from phage EPI vesicle to the nucleus stage. Combined, these systems seem to provide a multilayered defense to limit the escape possibilities of jumbo phages **(Figure 3G)**.

In summary, Ophion provides potent and specific defense against jumbo phages by blocking progression from the EPI vesicle to the nucleus stage, possibly by halting the transcription of the nucleus forming genes. The co-occurrence of Ophion with the EPI vesicle-targeting system Juk strongly suggests beneficial effects of targeting distinct features and processes of early jumbo phage infection.

### Ambrosia combines features of RM and two-component regulatory systems

Ambrosia, named after the food and drinks that provide immortality to the Greek gods, consists of five genes and provides protection against siphophages from the *Mesyanzhinovviridae* family and myophages from the *Pbunavirus* genus, limiting phage propagation in both solid and liquid cultures (**Figure 4A,B**). One step growth curve of øPa34 in control and Ambrosia cells revealed that the life cycle of the phage is severely compromised by the defense system, with a burst size significantly reduced from 165 ± 18 PFU/cell in control cells to undetectable in Ambrosia-expressing cells (**Figure 4C,D**). Ambrosia is found across Gammaproteobacteria, with the majority of its occurrences (76%) in Pseudomonadales (**Figure S3A; Table S2**).

**Figure 4.**
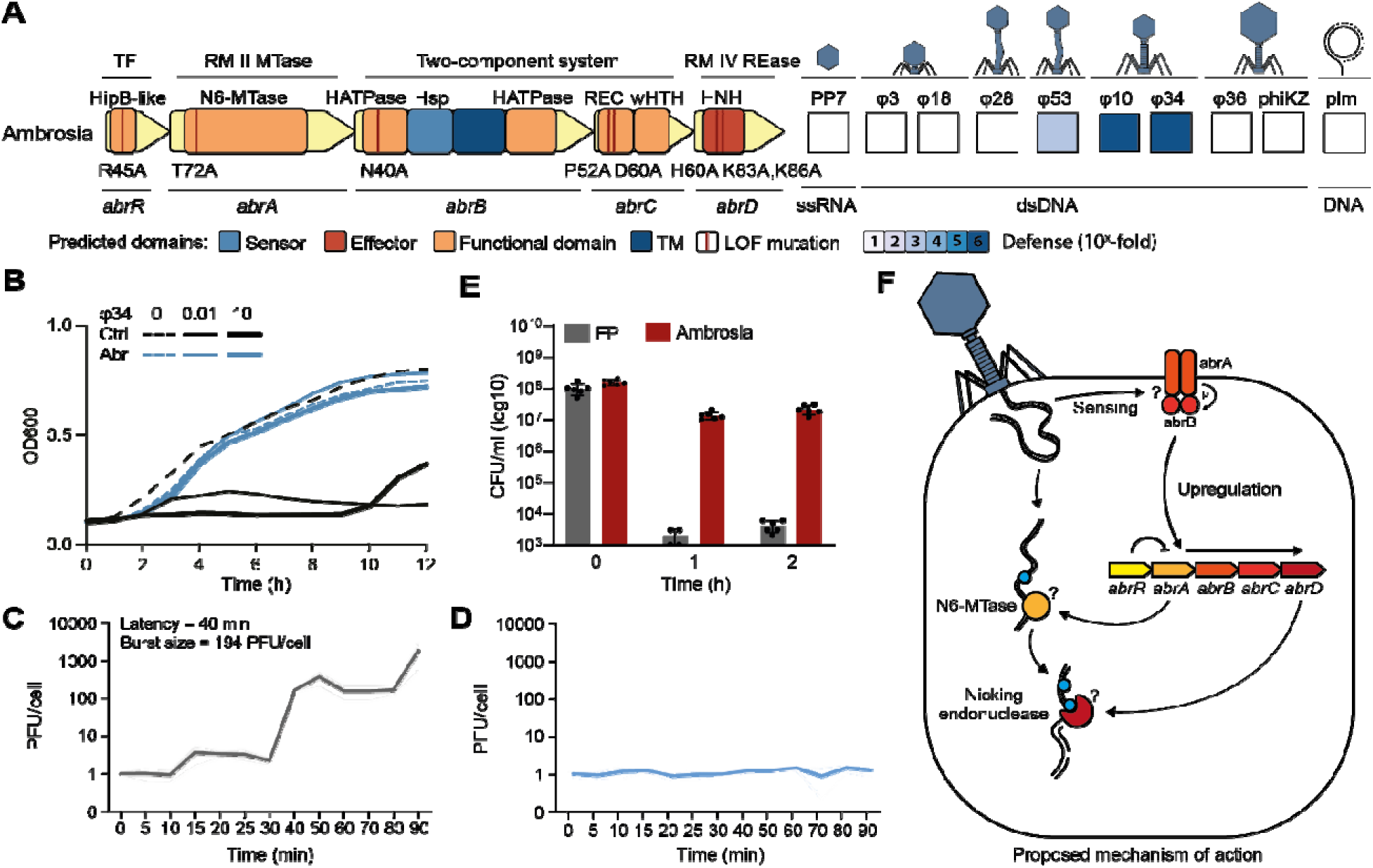
Ambrosia is a phage defense system that combines components of a restriction-modification system and a two-component system. **(A)** The functional domains of Ambrosia with the mutation sites tested indicated as red lines. Genes are drawn to scale and indicated for their association with phage defense (PD associated). Ambrosia was assessed for anti-plasmid activity (plm) by conjugation assays and for anti-phage activity by efficiency of plating assays. **(B)** Effect of Ambrosia on bacterial growth upon phage infection. *P. aeruginosa* strain PAO1 cells containing an empty plasmid (Ctrl), or Ambrosia (Abr) were infected with phage øPa34 at low (0.01) and high (10) multiplicity of infection, and their growth was monitored for a period of 12 h. **(C,D)** One-step growth curve of øPa34 infecting (C) control (empty plasmid) and (D) Ambrosia-expressing cells. **(E)** Assessment of cell concentration (CFU/ml) in phage-infected cultures of control (empty plasmid, EP) or Ambrosia-expressing cells. Bars represent average values of at least three biological replicates with individual points overlaid. **(F)** Working model for anti-phage defense by Ambrosia. Ambrosia is proposed to represent a defense system where a HipB-like XRE gene together with a two-component regulatory mechanism appears to control the expression of an RM-like system that targets its own DNA modification in response to phage infection.

The five genes of Ambrosia encode a HipB-like XRE transcriptional regulator (AbrR), a RM type II-associated N6 adenine-specific DNA methylase (MTase, AbrA), a sensor histidine kinase/heat-shock-protein(HSP)90-like ATPase (HATPase, AbrB), a response regulator (REC, AbrC), and an RM type IV HNH endonuclease (AbrD) (**Figure 4A; Figure S3B,C**). Each protein is required for anti-phage activity, since mutating conserved residues in any of the predicted functional domains completely abolishes Ambrosia defense activity (**Figure S3D**).

Ambrosia contains both elements from an RM system (MTase and HNH endonuclease^43^), which are known to be involved in phage defense, and a two-component regulatory system (HATPase and REC, a stimulus-response coupling mechanism^44^). Previous studies have reported that two-component systems often localize near RM systems, particularly RM types II and IV^45,46^, but their integration in a single gene cluster was not previously described. It has been proposed that the two-component system regulates the transcription of the RM systems, with the sensor kinase phosphorylating the response regulator to initiate transcription in response to specific environmental stimuli^47^. We hypothesized that this could also be the case for Ambrosia, especially since the REC-encoding AbrC contains a winged-helix-turn-helix (wHTH) domain, which is associated with these transcription initiating RECs^48^. In the context of Ambrosia, this suggests that AbrB detects phage infection and then phosphorylates AbrC, triggering a downstream transcriptional response to phage infection. The importance of AbrC phosphorylation is supported by the loss of phage protection when the predicted phosphorylated site (D60) was mutated to alanine (D60A) (**Figure S3D**). In addition, we quantified the expression of mNeonGreen inserted into the Ambrosia cluster in place of AbrA or AbrD, and observed that these regions were barely expressed in normal conditions but highly expressed during phage infection (**Figure S3E**). These results suggest that the expression of Ambrosia proteins is regulated at the transcription level.

In addition, Ambrosia expression may also be controlled by the HipB-like XRE transcriptional regulator AbrR, as these transcriptional regulators are known to inhibit transcription, especially of their own gene clusters^49^. In the context of RM systems, these XRE transcriptional repressors are known as control (C) proteins, which prevent the expression of the nuclease before the methylase has had time to methylate the host DNA^50^.

The apparent need for multiple transcriptional regulators to control Ambrosia gene expression suggests that its RM-like components could be toxic to the host when expressed in the absence of phage infection. Consistent with this idea, AbrA is predicted to be an N6 DNA methyltransferase and AbrD belongs to the RM type IV HNH endonuclease family, which typically cleaves methylated DNA. This raises the possibility that AbrD could act on modifications introduced by AbrA and, if expressed outside infection, damage host DNA. In classical type II RM systems, by contrast, the methylase protects host DNA from nuclease activity^51^.

To test whether AbrA methylation might provide protection from the nuclease of Ambrosia, we propagated phages in *P. aeruginosa* PAO1 expressing only AbrA from an inducible promoter and then challenged these phages against PAO1 carrying the complete Ambrosia system. In line with the hypothesis, phages remained sensitive, indicating that AbrA methylation does not protect against AbrD activity (**Figure S3F,G**). Interestingly, we did not observe toxicity of Ambrosia to the host during phage defense, which would have been expected if AbrD efficiently targeted the host chromosome (**Figure 4E**). One possibility is that AbrD activity is more damaging to phage DNA than to host DNA, as nicking nucleases may disproportionately affect linear genomes compared with circular chromosomes^52^.

Overall, our results indicate that the RM-like components of Ambrosia do not operate as a conventional type II RM system. Instead, Ambrosia appears to represent a distinct, tightly regulated phage defense system in which a HipB-like XRE protein and a two-component regulatory module control the expression of RM-like enzymes in response to infection (**Figure 4F**).

## Discussion

Evolutionary adaptation often proceeds from evolutionary tinkering processes, where existing genetic elements are repurposed or recombined^53^. Key to this process is the modular organization of genes and their constituent domains, which enables rapid exchange of distinct functional units among unrelated pathways in a “plug-and-play” fashion^53^. This is especially well exemplified by the core glycolytic genes, which are recruited across diverse metabolic pathways, including amino acid biosynthesis and aerobic respiration^54,55^.

Likewise, phage defense systems of bacteria exhibit high degrees of modularity, with sensing, signal transmission, and effector enzymes exchanged between phage defense systems^15^. This modularity may allow rapid adaptation to phage countermeasures^56–58^, for instance by replacing the sensory or effector domain that are inhibited by phage encoded anti-genes. In turn, the phage overcomes this adaptation, resulting in a continuous evolutionary race between the phage and the phage defense systems of the host.

In this study, we capitalized on the modularity of phage defense systems to discover three previously unknown defense systems in *P. aeruginosa* that incorporated genes or functional domains previously associated with phage defense in new architectures. Our discovery approach offers several powerful features. First, by searching for components already associated with phage defense, we increase the likelihood of finding positive candidate defense systems, independently of their abundance across bacterial genomes. Second, previously unknown anti-phage genes and functional domains can be integrated into subsequent searches, creating a feedback loop that continually expands our understanding of the phage defense repertoire. Lastly, this growing catalogue enables deeper insight into the evolutionary relationships between defense systems and the modular components they share. For example, many of the most frequently shared elements are RM components, and one of the systems uncovered by our approach, Ambrosia, combines an unusual RM-like module with regulatory genes.

The defense systems that we discovered using this approach include also Dionysus and Ophion. Dionysus contains a TerB-like and a pore-forming domain shared with several other defense systems but also contains an additional gene with a MAEBL functional domain that was not associated with phage defense before. Ophion features three genes with phage defense associated functionalities that have not previously been found in this combination. This system includes a rSAM, which is related to bacterial Viperins that convey phage defense^34,59^, along with PHP and PRTase domains. Both Dionysus and Ophion specifically block infection by jumbo phages, which have evolved specialized compartments to protect their DNA from DNA-targeting host defenses. After entry, the phage genome is enclosed by a membranous EPI vesicle, where early transcription occurs. At a later stage of the infection, the content of the EPI vesicle is transferred into a nucleus-like compartment where DNA replication and late transcription^32,60–63^. While these compartments provide jumbo phages with an advantage against some host defenses, our findings show that they can also be exploited as a vulnerability targeted by phage defenses such as Dionysus and Ophion.

Specifically, we show that Dionysus forms pores in the EPI vesicle of the jumbo phage to disrupt the infection cycle of the phage at an early stage, similar to the mechanism described for the Juk phage defense system^42^. In addition, we show that Ophion specifically prevents the progression of nucleus formation, potentially by halting phage transcription in the EPI vesicle stage. Together with jumbo phage specific Avs5 defense, which recognizes an early-expressed jumbo phage protein^64^, evidence is growing of the existence of a jumbo phage specific defense system repertoire. We also observed that Ophion almost exclusively co-localizes and co-occurs with the Juk phage defense system, suggesting evolutionary advantages of co-targeting.

Altogether, we explored the modularity of phage defense components across defense systems, expanding the known network of phage defense-associated domains employed by bacteria, and highlighting the diverse molecular functions recruited for phage defense.

## Data availability

Network files, phylogenetic trees, and HMM profiles of individual defense proteins have been deposited at Zenodo and are publicly available as of the data of publication at DOI: 10.5281/zenodo.17223413.

## Acknowledgments

This work was supported by the European Research Council CoG under the European Union’s Horizon 2020 research and innovation program (grant agreement no. 101003229 to S.J.J.B); and the Netherlands Organisation for Scientific Research (grant no. VI.C.192.027 to S.J.J.B.). A.M. is supported by grants from Koningin Wilhelmina Fonds (KWF) Unique High-Risk Project (grant agreement no. 15602) and NWO under Open Competition XS (grant agreement no. OCENW.XS23.1.006). We thank Dennis Claessen (Leiden University) for providing *E. coli* ET12567/pUZ8002 and Milan Gerovac (Helmholtz Centre for Infection Research) for providing phage PhiKZ. We acknowledge the infrastructure provided by Kavli Nanolab Imaging Centre for optical microscopy. We also thank members of the Brouns laboratory for the many discussions and ideas that improved our work.

## Author contributions

Conceptualization, S.J.J.B; Methodology, D.F.v.d.B. A.R.C., J.Q.E., A.M., H.v.d.B.; Formal Analysis, D.F.v.d.B. A.R.C., J.Q.E.; Investigation, D.F.v.d.B. A.R.C., J.Q.E., A.M., H.v.d.B.; Visualization, D.F.v.d.B. A.R.C., J.Q.E.; Writing – Original Draft, A.R.C., D.F.B.; Writing – Review & Editing, S.J.J.B.; Resources, S.J.J.B.; Funding Acquisition, A.M., S.J.J.B.

## Methods

### Bacterial strains and bacteriophages

The candidate defense systems were amplified from *P. aeruginosa* clinical isolates obtained from the University Medical Centre Utrecht^65^. pUCP20-based plasmids containing the defense systems were cloned in *Escherichia coli* strain Dh5α and subsequently in *P. aeruginosa* strain PAO1. *E. coli* strain ET12567/pUZ8002 was used in conjugation assays. All bacterial strains were grown in Lysogeny Broth (LB) at 37 °C and 180 rpm, or in LB agar (LBA, 1.5 % agar (w/v)) plates at 37 °C, unless stated otherwise. *E. coli* ET12567/pUZ8002 was grown in media supplemented with 50 µg/ml of kanamycin and 25 µg/ml of chloramphenicol. Strains containing plasmid pUCP20 were grown in media supplemented with 100 µg/ml of ampicillin (for *E. coli*) or 200 µg/ml of carbenicillin (for *P. aeruginosa*). Strains containing plasmid pSTDesR^66^ were grown in media supplemented with streptomycin at 25 µg/ml. *Pseudomonas* phage PP7 was acquired from LGC Standards, and all the remaining phages were obtained from the Fagenbank^65^. Phages were propagated in liquid media using *P. aeruginosa* strain PAO1 as the bacterial host. The resulting bacterial lysate was centrifuged at 3,000 × *g* for 15 min, filter-sterilized (0.2 µm PES), and stored at 4 °C until further use.

### Identification of putative anti-phage gene clusters in variable regions of *P. aeruginosa*

We used PPanGGOLiN v1.2.74^19,67^ to identify conserved gene clusters in the variable regions of all 541 complete *P. aeruginosa* assemblies from the RefSeq database as of June 16, 2022. Genes in these conserved gene clusters were analyzed for their proximity to one another using cblaster v1.3.19^68^, applying a threshold of 100-base-pair distance. Gene clusters encoding known and complete defense systems were identified using DefenseFinder v1.0.9 with defense-finder-models v1.2.2^69^. Additionally, the defense-finder models were used to identify incomplete phage defense systems. Lastly, anti-phage-related functional domains were identified in the sub-selected gene clusters using the Pfam-A^70^ and Superfamily HMM models^71^ using HMMer^72^ and Interproscan v5.60-92.0^73^ respectively, from a list of anti-phage-associated functional domains compiled from a literature search (**Table S3**). R package visNetwork (https://datastorm-open.github.io/visNetwork/) and Gephi^74^ were used to perform a network analysis of the multi-gene clusters that contained anti-phage associated functional domains in addition to other functional domains previously not associated with phage defense. Confirmed anti-phage gene clusters were further annotated using HHpred, InterProScan^73^, and CDD^75–77^. Moreover, each protein was also predicted using AlphaFold3^78,79^, searched for homologs with Foldseek^80^, and further analyzed for functional domains using CATH-AlphaFlow 2024^81^ domain predictions. Signal peptides and transmembrane regions were annotated using Phobius v1.01 and SignalP v6.0^82–84^.

### Cloning of candidate defense systems

The candidate defense systems were amplified from *P. aeruginosa* strains using Q5 DNA Polymerase (New England Biolabs) with primers from **Table S4**. The PCR products were run on 1% agarose gels, and bands were excised and cleaned using the Zymoclean Gel DNA Recovery Kit (Zymo Research). Plasmid pUCP20 was digested with BamHI and EcoRI (NEB), dephosphorylated with FastAP (Thermo Scientific), and cleaned with the Zymo DNA Clean & Concentrator Kit (Zymo Research). The defense systems were cloned into the digested pUCP20 using the NEBuilder HiFi DNA Assembly Master Mix (New England Biolabs) and transformed into chemically competent NEB^®^ 5-alpha Competent *E. coli* (New England Biolabs) following the manufacturer’s instructions. Plasmids (**Table S5**) were extracted using the GeneJET Plasmid Miniprep kit (Thermo Scientific) and confirmed by Sanger sequencing at Macrogen. Confirmed plasmids were transformed into *P. aeruginosa* strain PAO1 by electroporation as previously described ^85^ and the cells were plated on LBA plates supplemented with 200 µg/ml of carbenicillin.

### Selection and cloning of point mutations of the defense systems

Loss of function mutagenesis candidates were determined using a combination of literature search, multiple protein alignments using PSI-BLAST and ClustalW v2.1^86,87^, and Alphafold3 structural prediction^78,79^. Gene knockouts and point mutations of the defense systems in pUCP20 were obtained by round-the-horn site-directed mutagenesis using Q5 polymerase with the phosphorylated primers indicated in **Table S4**. The PCR products were digested with DpnI (New England Biolabs) to remove methylated template DNA and run on 1% agarose gels. The bands of the expected size were excised and cleaned with the Zymo Gel DNA Recovery Kit and ligated with T4 DNA ligase (New England Biolabs) at room temperature for 2 hours. The ligated products were transformed into chemically competent NEB^®^ 5-alpha Competent *E. coli* following the manufacturer’s instructions. Plasmids (**Table S5**) were extracted and sequenced as indicated above and transformed into *P. aeruginosa* strain PAO1 by electroporation.

### Efficiency of plating

Ten-fold serial dilutions of phage stocks were spotted onto double-layer agar (DLA) plates of *P. aeruginosa* strain PAO1 with empty pUCP20 or pUCP20 with the candidate defense systems following the small plaque drop assay^88^. The anti-phage activity of the systems was calculated as the fold reduction in phage infectivity of *P. aeruginosa* strain PAO1 that contains the defense system, compared to the phage infectivity of *P. aeruginosa* strain PAO1 containing the empty plasmid.

### Infection dynamics of phage-infected cultures

Overnight bacterial cultures of *P. aeruginosa* strain PAO1 with either empty or defense-containing pUCP20 were adjusted to an OD_600_ of 0.1 in LB and subjected to phage infection at an MOI lower than 1. The cultures were incubated at 37 °C and 180 rpm. Samples were collected at 0h, 2h, 4h, and 6h, followed by centrifugation at 3,000 × *g* for 5 min. The resulting phage-containing supernatant was serially diluted and spotted onto DLA plates of *P. aeruginosa* strain PAO1 to estimate phage concentration.

### Liquid culture collapse assays

Bacterial cultures grown overnight were diluted to an OD_600_ of 0.1 in LB and dispensed into 96-well plates. Phages were introduced at MOIs of 0.01 and 10, and the plates were incubated at 37 °C in an Epoch2 microplate spectrophotometer (Biotek). OD_600_ measurements were taken every 10 min over a 24h period, with double orbital shaking.

### Plasmid conjugation

Plasmid pSTDesR was conjugated into *P. aeruginosa* strain PAO1 containing empty pUCP20 or pUCP20 with individual defense systems using the puddle matting approach as previously described ^89^. Briefly, overnight cultures of the strains were diluted 1:2 in LB and incubated at 42°C for a minimum of 3h. Then, 0.5 ml of the *P. aeruginosa* cultures were mixed with 1.5 ml of an exponentially grown culture of *E. coli* ET12567/pUZ8002 containing pSTDesR. The cells were collected by centrifugation at 10,000 × *g* for 5 min, and resuspended in 50 μl of LB. The cell mixture was transferred onto the middle of a pre-warmed LBA plate and allowed to dry. The plates were incubated overnight at 30 °C. The bacterial puddle was then scaped off the surface of the LBA plate and resuspended in 1 ml of sterile phosphate buffer saline (1x PBS). Two-fold serial dilutions of this cell suspension were then platted on LBA plates containing either 100 μg/ml of carbenicillin (for total cell count) or additional 12.5 μg/ml streptomycin (for conjugants quantification). The plates were incubated for 48h at 37 °C.

### Adsorption assays

*P. aeruginosa* strain PAO1 carrying either an empty plasmid or a plasmid expressing a defense system was grown to early exponential phase (optical density at 600 nm [OD_600_] ≈ 0.3). Phages were added at a multiplicity of infection (MOI) of 0.01 and incubated at 37°C with shaking at 180 rpm for 10 minutes. Samples were collected at 0 and 10 minutes post-infection, centrifuged at 9,000LJ×LJ*g*, and the supernatant, containing non-adsorbed phages, was serially diluted 10-fold. The dilutions were plated on PAO1 double-layer agar plates to quantify non-adsorbed phage titers.

### One step growth curve

*P. aeruginosa* strain PAO1 carrying either an empty plasmid or a plasmid expressing a defense system was grown to an OD_600_ of 0.3–0.4. Cultures were centrifuged at 3,200LJ×LJ*g* for 10 minutes, and the cell pellet was resuspended in LB to half the original culture volume. Phages (øPa34 or øPa36) were added at a multiplicity of infection (MOI) of 0.01 and allowed to adsorb for 10 minutes at 37°C with shaking at 180 rpm. After adsorption, cultures were centrifuged, and the cell pellet was resuspended in LB to the original culture volume. Cultures were incubated at 37°C with shaking, for 90 minutes (Ambrosia) or 110 minutes (Ophion and Dionysus). For Ambrosia, samples were taken at time 0, every 5 minutes for the first 30 minutes, and then every 10 minutes thereafter. For Ophion and Dionysus, samples were collected at time 0 and at 10-minute intervals. Samples were immediately serially diluted 10-fold and spotted on PAO1 double-layer agar plates for phage quantification.

### Confocal microscopy

Confocal microscopy was performed as previously described^64^. *P. aeruginosa* strain PAO1 carrying either an empty plasmid or a plasmid expressing Dionysus or Ophion was grown to an OD_600_ of 0.3–0.4. Cells were infected with phage øPa36 at an MOI >3, and adsorption was allowed for 10 min at 37°C, 180 rpm. Cells were then centrifuged at 12,000 ×LJ*g* for 1 min to remove non-adsorbed phages, resuspended in LB, and incubated at 37°C, 180 rpm. Following incubation at different time points, cells were pelleted again and resuspended in 100 μl of LB. DAPI (to stain DNA) was added to final concentrations of 0.1 mg/ml. For cell viability staining, propidium iodide was added at a final concentration of 1 μM. Stained cells were spotted onto 1% agarose pads^90^ and imaged using a Nikon A1R/SIM laser scanning confocal microscope (inverted Nikon Ti Eclipse body) equipped with a 100× oil immersion objective (SR Apo TIRF; numerical aperture 1.49). DAPI was excited at 405 nm (emission filter: 450/50 nm), with signals passed through a 405/488/543/640 excitation dichroic mirror. Z-stacks were acquired using a Nikon A1 Piezo Z Drive at intervals of 0.1 or 0.2 µm (10–20 slices), capturing different focal planes of all bacteria in the field of view (512 × 512 pixels, corresponding to 36.79 × 36.79 µm, satisfying Nyquist criteria). A pinhole size corresponding to 1.2 Airy Units (AU), referenced to the shortest wavelength, was used. Images were acquired at 12-bit depth using a Galvano scanner with Nikon NIS-Elements software. Image analysis was performed using Fiji.

### Presence of the novel phage defense systems in bacteria

To detect all instances of Dionysus, Ophion, and Ambrosia in bacteria, we applied cblaster v1.3.18^68^ to obtain all clusters in the RefSeq database (minimum identity 20%). The taxonomy of all Dionysus, Ophion, and Ambrosia instances by cblaster was based on the annotation provided by RefSeq. Next, we built HMM profiles for the individual proteins of these defense systems using MUSCLE (-super5) v5.1^91^ to align the obtained protein sequences, and hmmbuild v3.3.2^72^ to build the HMM models. HMM models were searched against protein sequence databases using hmmsearch^72^. The HMM model scoring thresholds were set based on the 100% sensitivity point with the help of an ROC curve analysis that scored the HMM sensitivity for the defense protein compared to all other proteins within *P. aeruginosa*. The HMM bitscore obtained for each defense system protein is as follows: Dionysus: DinA, 400, DinB, 100 and DinC, 200; Ophion: OpnA, 500, OpnB, 350, and OpnC, 450; and Ambrosia: AbrA, 100, AbrB, 500, AbrC, 150, and AbrD, 200.

### Co-occurrence analysis

A pairwise co-occurrence analysis was performed using the presence and absence of all known jumbo phage defense systems, including Juk, Dionysus, CBASS type III, Avs5, 6A-MBL, and Ophion in the set of 541 complete *P. aeruginosa* assemblies from the RefSeq database as of June 16, 2022. The Z-scores and associated p-values were calculated based on the expected co-occurrence of these phage defense systems with the assumption of independence^92^. These p-values were then adjusted using the Bonferroni correction^93^.

### Phylogenetic analysis of shared phage defense genes

Phylogenetic trees of AbrA (N6-Methylase), AbrD (HNH), DinB (TerB), OpnB (PHP), and OpnC (PRTase), were made by obtaining all available Uniprot Release 2024_04^94^ proteins containing the corresponding functional domain. The phylogenetic tree of OpnA (rSAM) was built using the radical SAM containing protein list from Bernheim et al. (2021) ^34^. Duplicate sequences were removed, and the left-over proteins were subsequently downsampled, aligned, and trimmed as previously described^95^. The resulting alignments were used to build a phylogenetic tree using FastTree v2.1.11^96^ with default settings and visualized, without branch lengths, using the interactive Tree of Life (iTol) v6^97^. All previously known defense genes were then searched using the HMM profiles of Defense Finder Models v1.3.0 and PADLOC-DB v2.0.0^98,99^. A bitscore higher than 100 was seen as significant.

### Statistical analysis

Unless stated otherwise, data are presented as the means of biological triplicates ± standard deviation. A Bonferroni-adjusted p-value of less than 0.05 was considered significant.

## Supplemental information

**Supplementary file 1.** Geography markup language (.gml) format vector file of the modular gene network of phage defense systems, Related to **Figure 1**.

**Figure S1.** Characteristics of the phage defense system Dionysus, Related to Figure 2.

**Figure S2.** Characteristics of the phage defense system Ophion, Related to Figure 3.

**Figure S3.** Characteristics of the phage defense system Ambrosia, Related to Figure 4.

**Table S1.** Information on the conserved gene clusters found by PPanGGoLiN in the set of 541 RefSeq *Pseudomonas aeruginosa* genomes, related to STAR Methods.

**Table S2.** Identified instances of Dionysus, Ophion, and Ambrosia in bacteria, related to STAR Methods.

**Table S3.** Functional domains associated with phage defense, related to STAR Methods.

**Table S4.** List of primers used in this study, related to STAR Methods.

**Table S5.** List of plasmids used in this study, related to STAR Methods.

