## Supplementary for "Discovery of phage defense systems through component modularity networks"

### List of contents

**Figure S1.** Characteristics of the phage defense system Dionysus, Related to Figure 2.

**Figure S2.** Characteristics of the phage defense system Ophion, Related to Figure 3.

**Figure S3.** Characteristics of the phage defense system Ambrosia, Related to Figure 4.

**Table S4.** List of primers used in this study, Related to STAR Methods.

**Table S5.** List of plasmids used in this study, Related to STAR Methods.

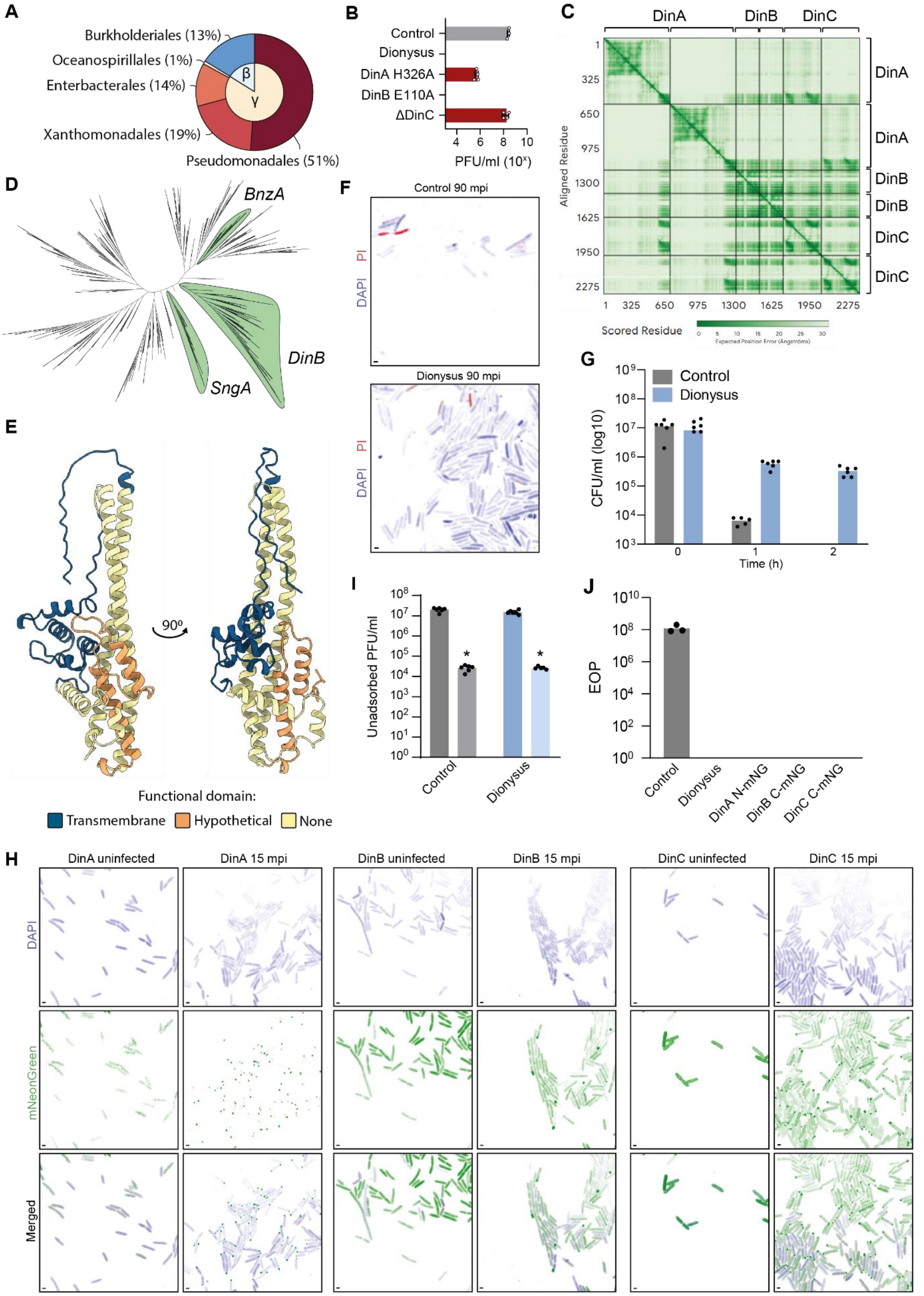


**Figure S1. Characteristics of the phage defense system Dionysus, Related to Figure 2.**

**(A)** The prevalence of Dionysus within Prokaryotes. Dionysus was found in a total of 713 instances, and its distribution (in percentage of total) in different orders is shown in the outer circles of the graphic, while the inner circle indicates the classes. γ: Gammaproteobacteria. β: Betaproteobacteria.

**(B)** Effect of gene deletions and mutations in the functional domains of Dionysus on phage protection. The infectivity of phage ϕPa36 on *P. aeruginosa* strain PAO1 cells containing an empty plasmid (control), Dionysus, or Dionysus with point mutations or deleted genes was assessed by plaque assay. Bars represent average values of at least three biological replicates with individual points overlaid.

**(C)** PAE plot obtained by co-folding the three Dionysus proteins (two copies of each) with AlphaFold3.

**(D)** Phylogenetic tree of TerB containing proteins. The phylogenetic tree of 1406 representative proteins were inferred using Fasttree2. The representative proteins include all groups within the TerB-like family (IPR029024), as indicated in the tree.

(**E**) Predicted Alphafold3 structure of alpha-PFT DinC.

**(F)** The membrane permeability of control and Dionysus-expressing cells 90 min post infection with ϕPa36, assessed with propidium iodide (PI) staining.

**(G)** The cell concentration (CFU/ml) of PAO1 containing an empty plasmid (control) or Dionysus, during ϕPa36 infection at an MOI of 10.

**(H)** Confocal microscopy images during ϕPa36 infection (0 and 15 min) of PAO1 expressing Dionysus with fluorescently labelled mNeonGreen-DinA, mNeonGreen-DinB, or mNeonGreen-DinC, stained with DAPI. Shown are the separate captured images for DAPI, and mNeonGreen, as well as the merged images.

**(I)** Adsorption assay of ϕPa36 on PAO1 containing an empty plasmid (control) or Dionysus.

**(J)** EOP assay of ϕPa36 in PAO1 containing an empty plasmid (control), Dionysus, or Dionysus with mNeonGreen (mNG) fused to N- or C-terminus of each Dionysus protein (DinA N-mNG, DinB C-mNG, and DinC C-mNG).

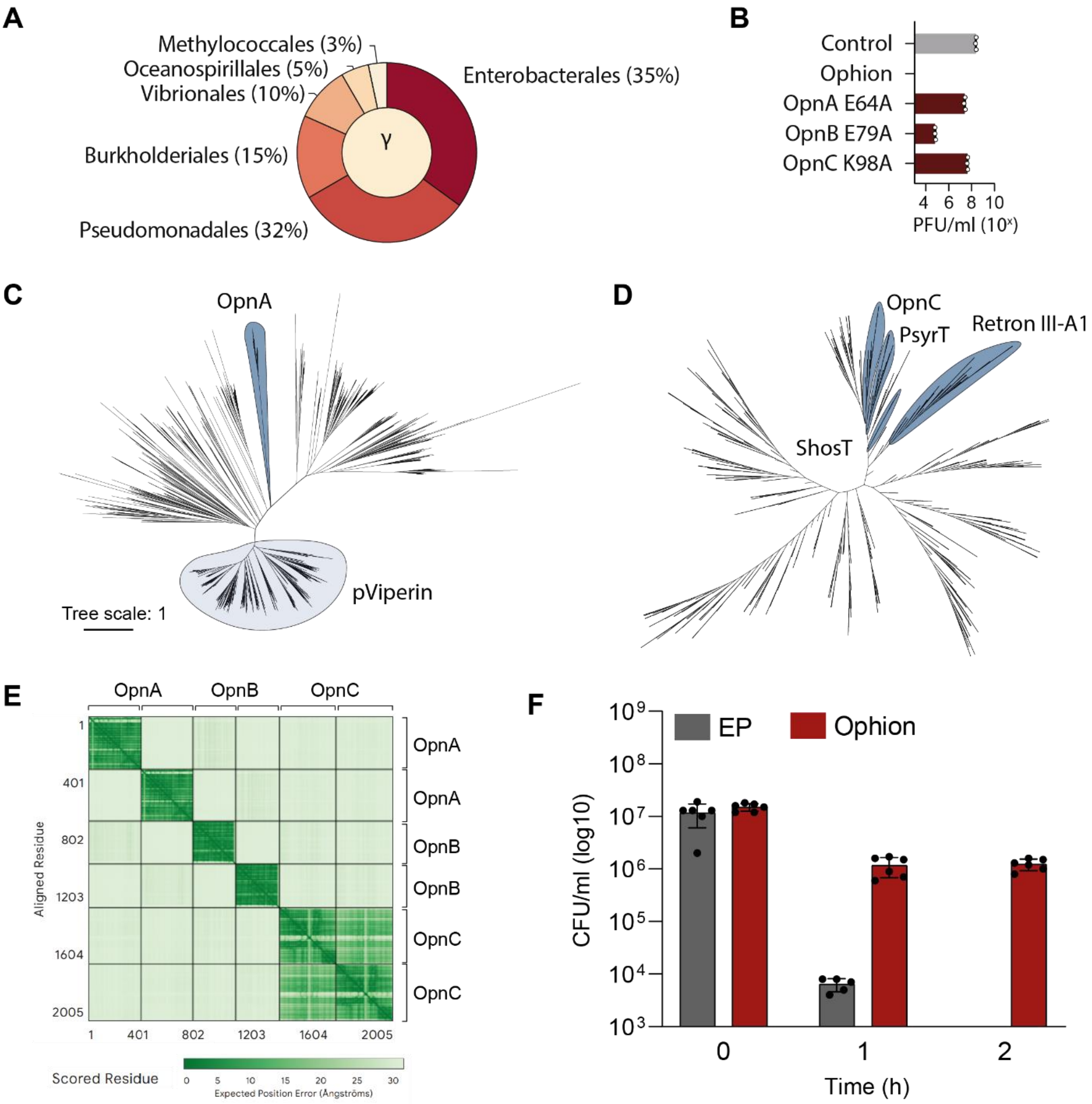


**Figure S2. Characteristics of the phage defense system Ophion, Related to Figure 3.**

**(A)** The prevalence of Ophion in prokaryotes. Ophion was found in a total of 4103 instances, and its distribution (in percentage of total) in different orders is shown in the outer circles of the graphic. γ: Gammaproteobacteria.

**(B)** Effect of mutations in the functional domains of Ophion on phage protection. The infectivity of phage ϕPa36 on *P. aeruginosa* strain PAO1 containing an empty plasmid (control), Ophion, or Ophion with point mutations was assessed by plaque assay. Bars represent average values of at least three biological replicates with individual points overlaid.

**(C)** Phylogenetic tree of radical SAM containing proteins. The phylogenetic tree of 1976 representative proteins was inferred using Fasttree2. The representative proteins include all groups within the radical SAM superfamily (PF04055), as indicated in the tree.

**(D)** Phylogenetic tree of PRTase-containing proteins. The phylogenetic tree of 1174 representative proteins was inferred using Fasttree2. The representative proteins include all groups within PFAM clan0533, as indicated in the tree.

**(E)** PAE plot obtained by co-folding all Ophion proteins (two copies each) using AlphaFold3.

**(F)** The cell concentration (CFU/ml) of PAO1 containing an empty plasmid (control) or Ophion, during infection of ϕPa36 at an MOI of 10.

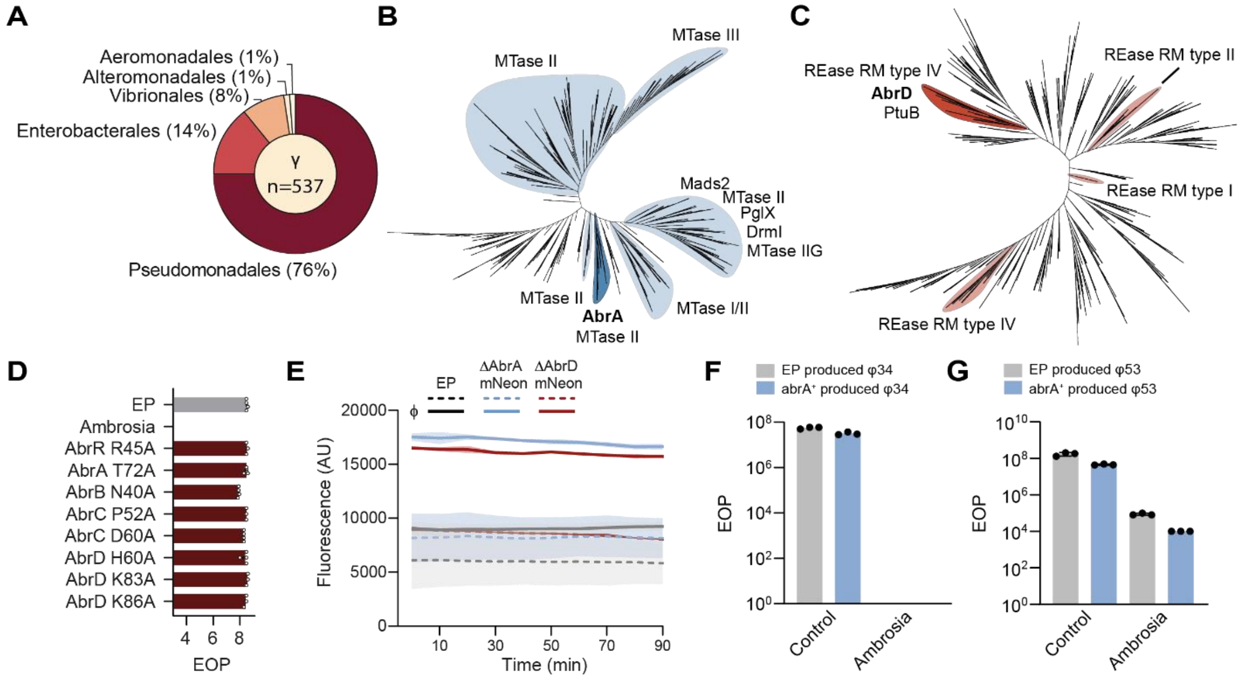


**Figure S3. Characteristics of the phage defense system Ambrosia, Related to Figure 4.**

**(A)** The prevalence of Ambrosia in prokaryotes. Ambrosia was found in a total of 537 instances, and its distribution (in percentage of total) in different orders is shown in the outer circles of the graphic. γ: Gammaproteobacteria.

**(B)** Phylogenetic tree of N6 MTase-containing proteins. The phylogenetic tree of 2066 representative proteins was inferred using Fasttree2. The representative proteins include all groups in N6-specific DNA methylases (IPR002052), as indicated in the tree.

**(C)** Phylogenetic tree of HNH containing proteins. The phylogenetic tree of 1681 representative proteins was inferred using Fasttree2. The representative proteins include all groups in HNH (IPR002711), as indicated in the tree.

**(D)** Effect of mutations in the functional domains of Ambrosia on phage protection. The infectivity of phage ϕPa34 on *P. aeruginosa* strain PAO1 cells containing an empty plasmid (control), Ambrosia, or Ambrosia with point mutations was assessed by plaque assay. Bars represent average values of at least three biological replicates with individual points overlaid.

**(E)** Fluorescence measurements (AU) over 90 minutes of infection by ϕPa34 on *P. aeruginosa* strain PAO1 cells containing an empty plasmid (EP), Ambrosia with *abrA* replaced by mNeonGreen, or Ambrosia with *abrD* replaced by mNeonGreen.

**(F)** EOP assay to assess the infectivity of ϕPa34 produced in PAO1 or PAO1:*abrA*$^{+}$, on PAO1 encoding an empty plasmid (control) or Ambrosia.

(**G**) EOP assay to assess the infectivity of ϕPa53 produced in PAO1 or PAO1 *abrA* $^{+}$, on PAO1 encoding an empty plasmid (control) or Ambrosia.

**Table S4.** List of primers used in this study, Related to STAR Methods

| Name | Sequence | Description |
|---|---|---|
| BN2830 | TGGTGAGACATGGGAAGCG | Sequencing primer pUCP20 |
| BN2840 | AGTGAGCGAGGAAGCGGAA | Sequencing primer pUCP20 |
| BN4448 | GCATGCCTGCAGGTCGACTCTAGAGGTGCCATG GTTCAGGTTC | Amplify Ambrosia from L0872 to clone into pUCP20 |
| BN4449 | GGAAACAGCTATGACCATGATTACGGATCAAGCCA CAACGAGAATAATAA | |
| BN4450 | GTTGAGTATCCATAAAAGGCTCCG | Sequencing primer Ambrosia |
| BN4451 | GTTCCACCGGAATGCTCTTC | Sequencing primer Ambrosia |
| BN4452 | GATTACGTCTTTATATGTGTTGAAGCT | Sequencing primer Ambrosia |
| BN4453 | ATGGTCTTTTAGCCCTGTCAGGAT | Sequencing primer Ambrosia |
| BN4454 | CTCGGGCAAGAAATCCACGG | Sequencing primer Ambrosia |
| BN4455 | GGGCACATTACTGGACTCGG | Sequencing primer Ambrosia |
| BN4531 | TGAATTTTCAGGAAATGCGGTGA | Sequencing primer pSTDesR |
| BN4532 | TCGGTCAAGGTTCATTTAAATACGT | Sequencing primer pSTDesR |
| BN4535 | TACAACAGGAAAGTAACGATGTCCGA | Sequencing primer Dionysus |
| BN4536 | ACGCCGCTGAGCCCTGAGC | Sequencing primer Dionysus |
| BN4537 | AGGAAAGAGCTACCAAACACCTCA | Sequencing primer Dionysus |
| BN4538 | GGCCATCAATATCTGTCTATCGCTGG | Sequencing primer Dionysus |
| BN4539 | CCGGAGCGCTTCTTGCCC | Sequencing primer Dionysus |
| BN4540 | GACTTCTCCACCACTGCGCT | Sequencing primer Dionysus |
| BN4578 | GCATGCCTGCAGGTCGACTCTAGAGTTTTATGTG TACGAACGATCTCGCACA | Amplify Dionysus from K6024 to clone into pUCP20 |
| BN4579 | GGAAACAGCTATGACCATGATTACGCTCGCTGAC GCCCATGCC | |
| BN4649 | P-GCGTCAAGTGCAAATCCTTCGTGTCC | Insert mutation H60A in AbrD from Ambrosia |
| BN4650 | P-TTTCCGCCCCCAAGAGCAAT | |
| BN4657 | P-GCAAGAATAATTGCTACTACACTTGG | Insert mutation D60A in AbrC from Ambrosia |
| BN4658 | P-CGAGCAGCTTAAAGAATCAG | |
| BN4663 | P-GCTTTAACGAGCTCGCTTATGG | Insert mutation N40A in AbrB from Ambrosia |
| BN4664 | P-CGCTTATGATGCTGATGCA | |
| BN4667 | P-CAAAAAAGACTCCTTGATGCTTTCG | Insert mutation T72A in AbrA from Ambrosia |
| BN4668 | P-CACCTGCCGAATTATCAAAATTAG | |
| BN4727 | GCATGCCTGCAGGTCGACTCTAGAGGACCT GCCCTGGGGCATTT | Amplify Ophion from L0872 to insert into pUCP20 |
| BN4728 | GGAAACAGCTATGACCATGATTACGAATTGC AGGTCAACGTCCCGTACT | |
| BN4739 | GATTGACGTGCTCCCGCTCC | Sequencing primer Ophion |
| BN4740 | GAAGGCCTGAGGCATTGAGC | Sequencing primer Ophion |
| BN4741 | CTAGGCTTCACCGTGACATC | Sequencing primer Ophion |
| BN4831 | P-GCGCCTTCGCCACTGACGG | Insert mutation E64A in OpnA of Ophion |
| BN4832 | P-GCCGCTCAACAACGCCAAAGTG | |
| BN4833 | P-GCCAGGCCGCACAGGATGG | Insert mutation E79A in OpnB of Ophion |
| BN4834 | P-ACTCGACTACATGCCTGGCACC | |
| BN4835 | P-GCTGCACGGCTGTAGATCCCG | Insert mutation K98A in OpnC of Ophion |
| BN4836 | P-ATACGGCAGTGTGGGTGATATCC | |
| BN4902 | P-GCGCTAGTCTTGAGGATGTTGGAGGCAG | Insert mutation H326A in DinA of Dionysus |
| BN4903 | P-CCGCGCCGGCATAGAGG | |
| BN5537 | TATCCTCCTCGCCCTTGCTCACCATTCTGCTTG | |

| | | |
|---|---|---|
| | CTCCGTCCTGGT | Replace AbrA in pUCP20-Ambrosia by mNeonGreen |
| BN5567 | GGGCATGGACGAGCTGTACAAGTAAATGGAAA TTAGAAAAAAAGATAGGCGATCACCCTTTCACTG | |
| BN5568 | TATCCTCCTCGCCCTTGCTCACCATTCATGATC TTTTAGAAAGTCCATTTTC | Replace AbrD in pUCP20-Ambrosia by mNeonGreen |
| BN5540 | GGGCATGGACGAGCTGTACAAGTAATTAACAAC TAGCACACTGCT | |
| BN5541 | P-GAGGTAGGTTCTCGCTTTCT | Deletion of DinC (ΔDinC) of Dionysus |
| BN5542 | P-AAGGAGAAATCCATGCCCAT | |
| BN5565 | CTGTTTCAAGCCCCTCAAATG | Sequencing the replacement of AbrA by mNeonGreen on Ambrosia |
| BN5566 | GTGCCATGGTTCAGGTTCG | Sequencing the replacement of AbrD by mNeonGreen on Ambrosia |
| BN5640 | AGAGGATCCCCGGGTACCGAGCTCGCTAATTT CCATCAGCCGCTC | Introduce methylase AbrA from Ambrosia into pUCP20 under inducible promoter |
| BN5641 | ACAGCTATGACCATGATTACGAATTTTGAGCATT AATGTATTAGAAGATCAGC | |
| BN5642 | CGCGAATTCGAGCTCGGTACCCGGGCTACA TCGGGTGGATTTCATTC | Introduce nuclease AbrD from Ambrosia into pSTDedR under inducible promoter |
| BN5643 | GGTATGCGCTCGACTCTAGAGGATCATGAAA CATCTATTTTTTCCCAGAATC | |
| BN5672 | P-GAGGTAGGTTCTCGCTTTCTC | Deletion of DinC (ΔDinB) from Dyonisus |
| BN5673 | P-AAGGAGAAATCCATGCCCATTC | |
| BN6034 | P-CGTCTTCGACTAAAAAACTCATATC | Insert mutation P52A in AbrC of Ambrosia |
| BN6035 | P-CAAGTGTAGTAGCAATTATTCTT | |
| BN6036 | P-GCGCTATTTAATGTTCGAAGATATG | Insert mutation K86A in AbrD of Ambrosia |
| BN6037 | P-AATCCCTCTATATATTCTTACC | |
| BN6309 | P-GCTCCACAGGCTATTATTAGATTATTG | Insert mutation K83A in AbrD of Ambrosia |
| BN6310 | P-ATGCAACAGAAACAAATCTTC | |
| BN6329 | P-GCTTCTTTTGATGTATAAATCAA | Deletion of AbrR (ΔAbrR) from Ambrosia |
| BN6330 | P-TTGAGCATTAATGTATTAGAAG | |
| BN6341 | CTTGTACAGCTCGTCCATGCC | Amplify mNeonGreen for N-terminal cloning |
| BN6342 | ATGGTGAGCAAGGGCGAGG | |
| BN6343 | TTACTTGTACAGCTCGTCCATGCC | Amplify mNeonGreen for C-terminal cloning |
| BN6344 | ATGGTGAGCAAGGGCGAGG | |
| BN6345 | TCCTCGCCCTTGCTCACCATGGATTTCTCC TTTTACTCTAGGAC | Amplify pUCP20-Dionysus to introduce mNeonGreem at DinC N-terminal |
| BN6346 | GCATGGACGAGCTGTACAAGATGCCCATTC CTCTCATCATC | |
| BN6347 | TCCTCGCCCTTGCTCACCATTGGAGTATCC AAGGAGTTTTTTTGG | Amplify pUCP20-Dionysus to introduce mNeonGreem at DinC C-terminal |
| BN6348 | TGGACGAGCTGTACAAGTAAAAGTGAAAGA AGGCCCGTCAG | |
| BN6349 | TCCTCGCCCTTGCTCACCATGAGGTAGGTT CTCGCTTTCTCG | Amplify pUCP20-Dionysus to introduce mNeonGreem at DinB N-terminal |
| BN6350 | GCATGGACGAGCTGTACAAGATGCTCATCA AGATGCTGCC | |
| BN6351 | TCCTCGCCCTTGCTCACCATCTCTAGGACG AGGGCTATGG | Amplify pUCP20-Dionysus to introduce mNeonGreem at DinB C-terminal |
| BN6352 | TGGACGAGCTGTACAAGTAAAAGGAGAAAT CCATGCCCATTC | |
| BN6353 | TCCTCGCCCTTGCTCACCATGGGAGTTCCT GAGAAAGTGTGGTAAG | Amplify pUCP20-Dionysus to introduce mNeonGreem at DinA N-terminal |
| BN6354 | GCATGGACGAGCTGTACAAGATGAAGCCCC ATGCGCCCTC | |
| BN6355 | TCCTCGCCCTTGCTCACCATCAGGATCAGT GGCTGGTCGG | Amplify pUCP20-Dionysus to introduce mNeonGreem at DinA C-terminal |
| BN6356 | TGGACGAGCTGTACAAGTAACGAGAAAGCG AGAACCTACCTC | |
| BN6365 | P-GCCTCAAATCTGGATAGAGACTCTTGGGT TATG | Insert mutation R45A in AbrR from Ambrosia |
| BN6366 | P-GGGCCGTGCTGCTGAGTTTG | |

**Table S5.** List of plasmids used in this study, Related to STAR Methods

| Name in this study | Code | Insert | Derived from vector | Resistance marker |
|---|---|---|---|---|
| pUCP20 | pTU646 | - | - | $Amp^R$ |
| pUCP20-Dionysus | pTU1134 | Dionysus system from K6024 | pUCP20 | $Amp^R$ |
| pUCP20-Ophion | pTU1135 | Ophion system from L0872 | pUCP20 | $Amp^R$ |
| pUCP20-Ambrosia | pTU1136 | Ambrosia system from L0872 | pUCP20 | $Amp^R$ |
| pUCP20-Dionysus-H326A | pTU1137 | H326A mutation in DinA | pUCP20-Dionysus | $Amp^R$ |
| pUCP20-Dionysus-ΔDinB | pTU1138 | ΔDinB | pUCP20-Dionysus | $Amp^R$ |
| pUCP20-Dionysus-ΔDinC | pTU1139 | ΔDinC | pUCP20-Dionysus | $Amp^R$ |
| pUCP20-Ophion-E64A | pTU1140 | E64A mutation in OpnA | pUCP20-Ophion | $Amp^R$ |
| pUCP20-Ophion-E79A | pTU1141 | E79A mutation in OpnB | pUCP20-Ophion | $Amp^R$ |
| pUCP20-Ophion-K98A | pTU1142 | K98A mutation in OpnC | pUCP20-Ophion | $Amp^R$ |
| pUCP20-Ambrosia-ΔAbrR | pTU1143 | ΔAbrR | pUCP20-Ambrosia | $Amp^R$ |
| pUCP20-Ambrosia-E45A | pTU1144 | E45A mutation in AbrR | pUCP20-Ambrosia | $Amp^R$ |
| pUCP20-Ambrosia-T72A | pTU1145 | T72A mutation in AbrA | pUCP20-Ambrosia | $Amp^R$ |
| pUCP20-Ambrosia-N40A | pTU1146 | N40A mutation in AbrB | pUCP20-Ambrosia | $Amp^R$ |
| pUCP20-Ambrosia-P52A | pTU1147 | P52A mutation in AbrC | pUCP20-Ambrosia | $Amp^R$ |
| pUCP20-Ambrosia-D60A | pTU1148 | D60A mutation in AbrC | pUCP20-Ambrosia | $Amp^R$ |
| pUCP20-Ambrosia-H60A | pTU1149 | H60A mutation in AbrD | pUCP20-Ambrosia | $Amp^R$ |
| pUCP20-Ambrosia-K83A | pTU1150 | K83A mutation in AbrD | pUCP20-Ambrosia | $Amp^R$ |
| pUCP20-Ambrosia-K86A | pTU1151 | K86A mutation in AbrD | pUCP20-Ambrosia | $Amp^R$ |
| pUCP20-Dionysus-DinA-mNeonGreen | pTU1152 | mNeonGreen fused to C-terminal of DinA | pUCP20-Dionysus | $Amp^R$ |
| pUCP20-Dionysus-DinB-mNeonGreen | pTU1153 | mNeonGreen fused to C-terminal of DinB | pUCP20-Dionysus | $Amp^R$ |
| pUCP20-Dionysus-DinC-mNeonGreen | pTU1154 | mNeonGreen fused to C-terminal of DinC | pUCP20-Dionysus | $Amp^R$ |
| pUCP20-Ophion-OpnA-mNeonGreen | pTU1155 | mNeonGreen fused to C-terminal of OpnA | pUCP20-Ophion | $Amp^R$ |
| pUCP20-Ophion-OpnB-mNeonGreen | pTU1156 | mNeonGreen fused to C-terminal of OpnB | pUCP20-Ophion | $Amp^R$ |
| pUCP20-Ophion-OpnC-mNeonGreen | pTU1157 | mNeonGreen fused to C-terminal of OpnC | pUCP20-Ophion | $Amp^R$ |
| pUCP20-Ambrosia-ΔAbrA:mNeonGreen | pTU1158 | AbrA replaced by mNeonGreen | pUCP20-Ambrosia | $Amp^R$ |
| pUCP20-Ambrosia-ΔAbrD:mNeonGreen | pTU1159 | AbrD replaced by mNeonGreen | pUCP20-Ambrosia | $Amp^R$ |

| | | | | |
|---|---|---|---|---|
| pUCP20-AbrA | pTU1160 | AbrA under inducible promoter | pUCP20 | $Amp^R$ |
| pSTDesR | pTU679 | - | - | $Strep/Spec^R$ |
| pSTDesR-AbrD | pTU1161 | AbrD under inducible promoter | pSTDesR | $Strep/Spec^R$ |
